# Quantitative Evaluation of Nonlinear Methods for Population Structure Visualization & Inference

**DOI:** 10.1101/2022.02.22.481549

**Authors:** Jordan Ubbens, Mitchell J. Feldmann, Ian Stavness, Andrew G. Sharpe

## Abstract

Population structure (also called genetic structure and population stratification) is the presence of a systematic difference in allele frequencies between sub-populations in a population as a result of non-random mating between individuals. It can be informative of genetic ancestry, and in the context of medical genetics it is an important confounding variable in genome wide association studies. Recently, many nonlinear dimensionality reduction techniques have been proposed for the population structure visualization task. However, an objective comparison of these techniques has so far been missing from the literature. In this paper, we discuss the previously proposed nonlinear techniques and some of their potential weaknesses. We then propose a novel quantitative evaluation methodology for comparing these nonlinear techniques, based on populations for which pedigree is either known *a-priori* through artificial selection or simulation. Based on this evaluation metric, we find graph-based algorithms such as t-SNE and UMAP to be superior to PCA, while neural network based methods fall behind.

## Introduction

Population structure—the patterns of ancestral similarities and dissimilarities with arbitrarily defined populations—is a topic of primary concern in population, quantitative, and evolutionary genetics in humans, plants, microbes, and animals. The basic cause of population structure in sexually reproducing species is non-random mating between groups: if all individuals within a population mate randomly, then the allele frequencies should be similar between groups. Population structure commonly arises from physical or reproductive separation and isolation by distance, barriers, like mountains and rivers, followed by genetic drift (Holsinger and Weir, 2009; Petkova et al., 2016). Other causes include gene flow from migrations, population bottlenecks and expansions, founder effects, evolutionary pressure, random chance, and (in humans) cultural factors. As such, it should be expected that clusters, such as families, tribes, and clades should appear naturally in the data (Battey et al., 2021; Holsinger and Weir, 2009).

Estimates of population structure are usually derived from linear factor models such as Principal Components Analysis (PCA), but nonlinear dimensionality reduction techniques have attracted interest in recent literature. Dimensionality reduction has been an important tool for geneticists and has been widely used both to control for the effects of population structure in GWAS (Patterson et al., 2006; Price et al., 2006; Yu et al., 2006) and visualization and inference of genetic variation, i.e., population structure (Battey et al., 2021; Marnetto and Huerta-Sánchez, 2017; Steinig et al., 2016; Francis, 2017).

One existing challenge with estimating population structure is that there is no ground truth with which to compare. Sometimes, geographic distances are used as correlates of euclidean distance in principal component space to measure population reconstruction and representation (Battey et al., 2021). Given patterns of limited dispersal, on average, in many natural species it would make sense the genetic distance would be correlated with measures of geographic distance (Battey et al., 2021; Van Heerwaarden et al., 2011). However, for many locally restricted species and populations, e.g., breeding programs, geography is not a correct ground truth metric by which to judge tools for population structure visualization and inference.

Here, using pedigreed populations as well as simulations, we deploy a ground-truthing method for comparing and contrasting dimensionality reduction (embedding) techniques for population structure visualization and inference. We simulate populations with or without assortative mating (selection) and with or without migration (population structure) and keep track of the pedigree, sub-population membership, and individual level genotypes over multiple generations. We track the length of the shortest path between individuals in the pedigree network as the ground truth metric for ancestral distance. Pairs of individuals across more distantly related sub-populations will have a greater number of edges between them, compared to pairs of individuals within subpopulations, because they share a more distant most recent common ancestor. We use this ground truth to compare PCA, t-SNE, UMAP, an autoencoder, a variational autoencoder, constrastive embedding learning, and random projections. In these experiments, UMAP and t-SNE significantly outperform all other methods regardless of mating, selection, or structure.

### Review of Existing Nonlinear Methods

There are several nonlinear models for population structure visualization and inference which have been demonstrated in the literature. These methods can generally be divided into three categories: graph-based algorithms, autoencoders (AE), and variational autoencoders (VAEs). Here, we briefly discuss these algorithms and their applications.

#### Graph-Based Algorithms

Two of the most popular techniques used for visualizing high-dimensional data, which have both seen application in population genetic data, are t-SNE (Platzer, 2013; Li et al., 2017) and UMAP (Diaz-Papkovich et al., 2019). These methods are part of a family of techniques which are concerned with representing the data as a graph with various edge lengths. These techniques

#### Autoencoders

An autoencoder is a type of neural network which is commonly used for learning a set of features from data in an unsupervised fashion. Recently, autoencoders have been applied to population genetic data (Ausmees and Nettelblad, 2020; López-Cortés et al., 2020). Autoencoders are comprised of two components, an encoder neural network and a decoder neural network, which jointly reconstruct an input:

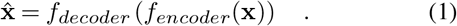

The encoder network *f_encoder_* compresses the highdimensional input **x** into a lower-dimensional space, while the decoder network *f_decoder_* outputs an approximation 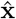 of the original input using only the low-dimensional embedding provided by the encoder. Since the embedding space acts as a bottleneck, it must efficiently represent as much of the relevant information as possible from the original signal. For this reason, autoencoders have often been used as a means of extracting informative and discriminative features for downstream tasks, where the embedding representation can be reused for another purpose. Specialized autoencoders can also be used for tasks such as denoising input data, like images (Vincent et al., 2008).

#### Variational Autoencoders

The variational autoencoder, or VAE, is a latent variable model which defines a generative model over data (Kingma and Welling, 2013). VAEs have been used for population data using both the standard Gaussian prior (Battey et al., 2021), as well as a Gaussian Mixture prior (GMM-VAE) (Meisner and Albrechtsen, 2020). When learning a latent variable model such as a VAE, one would like to maximize the probability of the data under the model

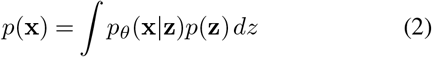

where **z** is the latent variable and *θ* are the model parameters. In this case, performing exact inference is analytically intractable due to the integral in Equation 2. However, we are able to minimize the *evidence lower bound* (ELBO) directly, given by

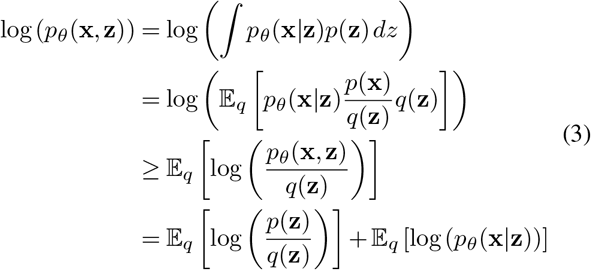

where *q* is the tractable prior. VAEs are trained using stochastic sampling, which we will skip here for brevity, and is described in more detail in Kingma and Welling (2013). VAEs and their many variants are useful models because the posterior density of the model approximates the tractable prior. This means that sampling new data from the complex, multimodal data distribution is as simple as sampling from the known prior. It also means that, unlike with the autoencoder, the density of the posterior is approximately continuous in the latent space and so it is often possible to smoothly interpolate between points in this space.

### Challenges with Existing Nonlinear Methods

#### Methods with Nonlinear Decoders Distort Distances Arbitrarily

Autoencoders, VAEs, and variants use decoders which are parameterized by neural networks. It is well-established that, when the decoder is a nonlinear function, the Euclidean distance between points in the embedding space is meaningless, as these distances are distorted by the decoder (Arvanitidis et al., 2017). The amount of instantaneous distortion is given by the Jacobian of the decoder evaluated with respect to **z**. This is straightforward to intuit: imagine moving through the embedding space while monitoring the output of the decoder. There are areas of the space where moving a short distance in a straight line elicits large changes in the output of decoder, and other areas where moving even a large distance does not change the output of the decoder at all. This poses a problem when using these methods for visualization and inference, as it is natural to interpret distance between points in the visualization in terms of Euclidean distance, without knowing that the data actually lie on some nonlinear sub-manifold embedded in the space. It is important to note that just because a nonlinear decoder is absent, does not mean that Euclidean distance is always respected – some methods, such as t-SNE, prefer nonlinear (e.g. t-distributed) distance for a specific visualization purpose, for example to emphasize local differences and avoid the over-crowding of points in a local area.

#### VAEs are Sensitive to the Choice of Prior

The VAE is a powerful generative model. However, the inference model *q_θ_*(**z**|**x**) learned by the VAE is generally meaningless with respect to the true data generating distribution. The ELBO is only a lower bound on the model evidence, and the difference between the lower bound and the true model evidence is given by

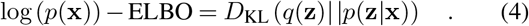

Therefore, the model is an unboundedly bad approximation of the true data distribution, where the quality of that approximation is given by the Kullback-Leibler divergence between *q* and the true distribution. Simply put, the samples will be approximately distributed according to whatever prior is chosen, regardless of what the actual underlying data generating distribution is. Using an implicit prior, one could even make the samples take on arbitrary distributions (Huszár, 2017). For this reason, if the model will be used for making inferences about population structure, then the user must ensure that the chosen prior accurately represents the true underlying distribution, making the lower bound tight.

#### Methods Explicitly Modelling Clusters May Result in Spurious Clustering

Population stratification naturally results in uneven distribution of alleles, but not necessarily completely discrete, compact clusters. For example, gradual migration over time may result in more of a gradient of allele frequency than in *K* completely distinct groupings. Methods which assign individuals to clusters, such as GMM-VAE, may exaggerate the presence of clusters, when the actual distribution of alleles is less discrete.

## Results & Discussion

For all experiments, we report Pearson’s correlation between the point-to-point distance in the visualization and the known ancestral distance (as given by the pedigree records or the simulation). We provide two sets of results, based on two potential interpretations of how distance between ancestors should be portrayed visually. The upper portion of Table 1 shows results where the simple Euclidean distance between points in the visualization is used, while the lower portion of Table 1 shows results where log_2_ of this distance is used. The former is based on the assumption that pedigree distance should be represented linearly, independent of depth, while the latter assumes that distance between points should correlate with the proportion of ancestry represented by that ancestor. For example, if an individual is separated from their parent by a distance of four units, then the linear distance assumes that their grandparent should be eight (2×4) units away, while the exponential distance assumes that the grandparent should be sixteen (4^2^) units away. We found that, in general, the log_2_ definition resulted in higher correlations with marker inferred distance.

**Table 1.** Correlation between the Euclidean distance, *D*, and log_2_ of the Euclidean distance, log_2_(*D*), in the visualization and the known pedigree distance. For the simulations, the superscript letters are the compact letter display for the Tukey least significant difference test (shown in Figure 3).

| Distance Metric | Model | Strawberry | Pig | Soay Sheep | Florida Scrub-Jay | Migration |  | No Migration |  |
| --- | --- | --- | --- | --- | --- | --- | --- | --- | --- |
|  |  |  |  |  |  | Random | Assortative | Random | Assortative |
| $D$ | UMAP | 0.85 | 0.73 | **0.78** | **0.18** | **0.80** ($\pm 0.19$)<sup>e</sup> | **0.78** ($\pm 0.17$)<sup>d</sup> | **0.16** ($\pm 0.02$)<sup>e</sup> | **0.24** ($\pm 0.03$)<sup>d</sup> |
| | t-SNE | 0.76 | **0.77** | 0.75 | 0.17 | 0.72 ($\pm 0.20$)<sup>d</sup> | 0.74 ($\pm 0.18$)<sup>d</sup> | 0.11 ($\pm 0.01$)<sup>d</sup> | 0.19 ($\pm 0.03$)<sup>d</sup> |
| | PCA | **0.86** | 0.48 | 0.07 | 0.15 | 0.76 ($\pm 0.21$)<sup>d,e</sup> | 0.75 ($\pm 0.20$)<sup>d</sup> | 0.11 ($\pm 0.02$)<sup>d</sup> | 0.18 ($\pm 0.04$)<sup>d</sup> |
| | β-VAE | 0.57 | 0.41 | 0.14 | 0.14 | 0.51 ($\pm 0.18$)<sup>c</sup> | 0.51 ($\pm 0.17$)<sup>c</sup> | 0.05 ($\pm 0.02$)<sup>c</sup> | 0.11 ($\pm 0.03$)<sup>c</sup> |
| | contrastive | 0.73 | 0.54 | 0.12 | 0.16 | 0.49 ($\pm 0.21$)<sup>c</sup> | 0.52 ($\pm 0.22$)<sup>c</sup> | 0.05 ($\pm 0.01$)<sup>c</sup> | 0.11 ($\pm 0.03$)<sup>c</sup> |
| | autoencoder | 0.76 | 0.44 | 0.01 | 0.14 | 0.35 ($\pm 0.16$)<sup>b</sup> | 0.35 ($\pm 0.18$)<sup>b</sup> | 0.03 ($\pm 0.01$)<sup>b</sup> | 0.05 ($\pm 0.03$)<sup>b</sup> |
| | random | 0.06 | 0.07 | 0.11 | 0.10 | 0.04 ($\pm 0.03$)<sup>a</sup> | 0.07 ($\pm 0.05$)<sup>a</sup> | 0.02 ($\pm 0.01$)<sup>a</sup> | 0.03 ($\pm 0.01$)<sup>a</sup> |
| $\log_2(D)$ | UMAP | 0.71 | **0.72** | **0.70** | 0.29 | **0.85** ($\pm 0.16$)<sup>e</sup> | **0.82** ($\pm 0.12$)<sup>e</sup> | **0.28** ($\pm 0.02$)<sup>f</sup> | **0.39** ($\pm 0.03$)<sup>f</sup> |
| | t-SNE | 0.64 | 0.70 | 0.64 | **0.31** | 0.75 ($\pm 0.19$)<sup>d</sup> | 0.78 ($\pm 0.15$)<sup>d,e</sup> | 0.19 ($\pm 0.02$)<sup>e</sup> | 0.30 ($\pm 0.03$)<sup>e</sup> |
| | PCA | **0.74** | 0.57 | 0.14 | 0.16 | 0.74 ($\pm 0.20$)<sup>d</sup> | 0.74 ($\pm 0.19$)<sup>d</sup> | 0.10 ($\pm 0.02$)<sup>d</sup> | 0.17 ($\pm 0.04$)<sup>d</sup> |
| | β-VAE | 0.53 | 0.51 | 0.22 | 0.16 | 0.62 ($\pm 0.20$)<sup>c</sup> | 0.62 ($\pm 0.17$)<sup>c</sup> | 0.06 ($\pm 0.02$)<sup>b</sup> | 0.15 ($\pm 0.03$)<sup>c</sup> |
| | contrastive | 0.57 | 0.56 | 0.12 | 0.17 | 0.50 ($\pm 0.21$)<sup>b</sup> | 0.52 ($\pm 0.21$)<sup>b</sup> | 0.05 ($\pm 0.01$)<sup>b</sup> | 0.11 ($\pm 0.03$)<sup>b</sup> |
| | autoencoder | 0.50 | 0.33 | 0.21 | 0.15 | 0.52 ($\pm 0.17$)<sup>b</sup> | 0.48 ($\pm 0.14$)<sup>b</sup> | 0.07 ($\pm 0.02$)<sup>c</sup> | 0.11 ($\pm 0.03$)<sup>b</sup> |
| | random | 0.22 | 0.10 | 0.12 | 0.07 | 0.03 ($\pm 0.02$)<sup>a</sup> | 0.07 ($\pm 0.05$)<sup>a</sup> | 0.02 ($\pm 0.01$)<sup>a</sup> | 0.03 ($\pm 0.01$)<sup>a</sup> |

### Comparison of Linear & Nonlinear Methods

Our evaluations show that nonlinear methods are often superior alternatives to PCA for visualizing population structure. t-SNE or UMAP outperformed PCA in every dataset, with the exception of strawberry. The strength of PCA in the strawberry example may be due to ascertainment bias in genotyping arrays, depth and complexity of the pedigree records, strength of selection between different sup-groups, or the over-representation of modern elite *F*. × *ananassa* in the database. PCA was only outperformed by one or more of the neural network methods (autoencoder, *β*-VAE, and contrastive) in three of the eight datasets when linear distance was assumed, and two of the eight when exponential distance was assumed, demonstrating that there is little incentive to use these techniques over PCA as a standard choice. However, these methods did consistently perform better than the random projection, indicating that they do capture significant information about ancestry.

In addition to the quantitative results, the visualizations in Figure 1C of the simulated population in Figure 1B illustrate some key differences between the methods. The groundtruth plot shows that there are five major clusters of subpopulations, with each one broken down further into multiple smaller subpopulations. Only some of these clusters are captured with PCA, as the first two principal components are insufficient to differentiate between some of the more closely related subpopulations. The autoencoder shows its characteristic “smearing” of points in the space, while clusters are vaguely visible. It is apparent that the autoencoder does not consider pairwise Euclidean distance to be important – the heads of each cluster are closer to each other than to the points in their tails. The unit Gaussian prior used in the *β*-VAE pushes the bulk of the clusters together towards the mode of the prior distribution at the origin. The distribution of points is somewhat removed from the prior, due to the relaxed KL term in the *β*-VAE. Meanwhile, the use of cosine distance in the contrastive method creates a ring-like structure, as the positions of the points are based on the angle they make with the origin. t-SNE and UMAP both show the major clusters clearly, although UMAP clusters individuals more tightly within each group, while t-SNE prefers to spread each cluster so individual points can be seen more clearly. This difference may account for the advantage UMAP holds over t-SNE in the quantitative results. Hyperparameters could also be tuned to decrease the tightness of the clusters in UMAP, if larger clusters are preferable for visualization reasons. Compared to the other techniques, the advantages of t-SNE and UMAP are clearly demonstrated here. These methods do not lose important information like PCA, which collapses several of the subpopulations together, and they respect pairwise Euclidean distances between individuals better than the neural network-based methods.

**Figure 1.**
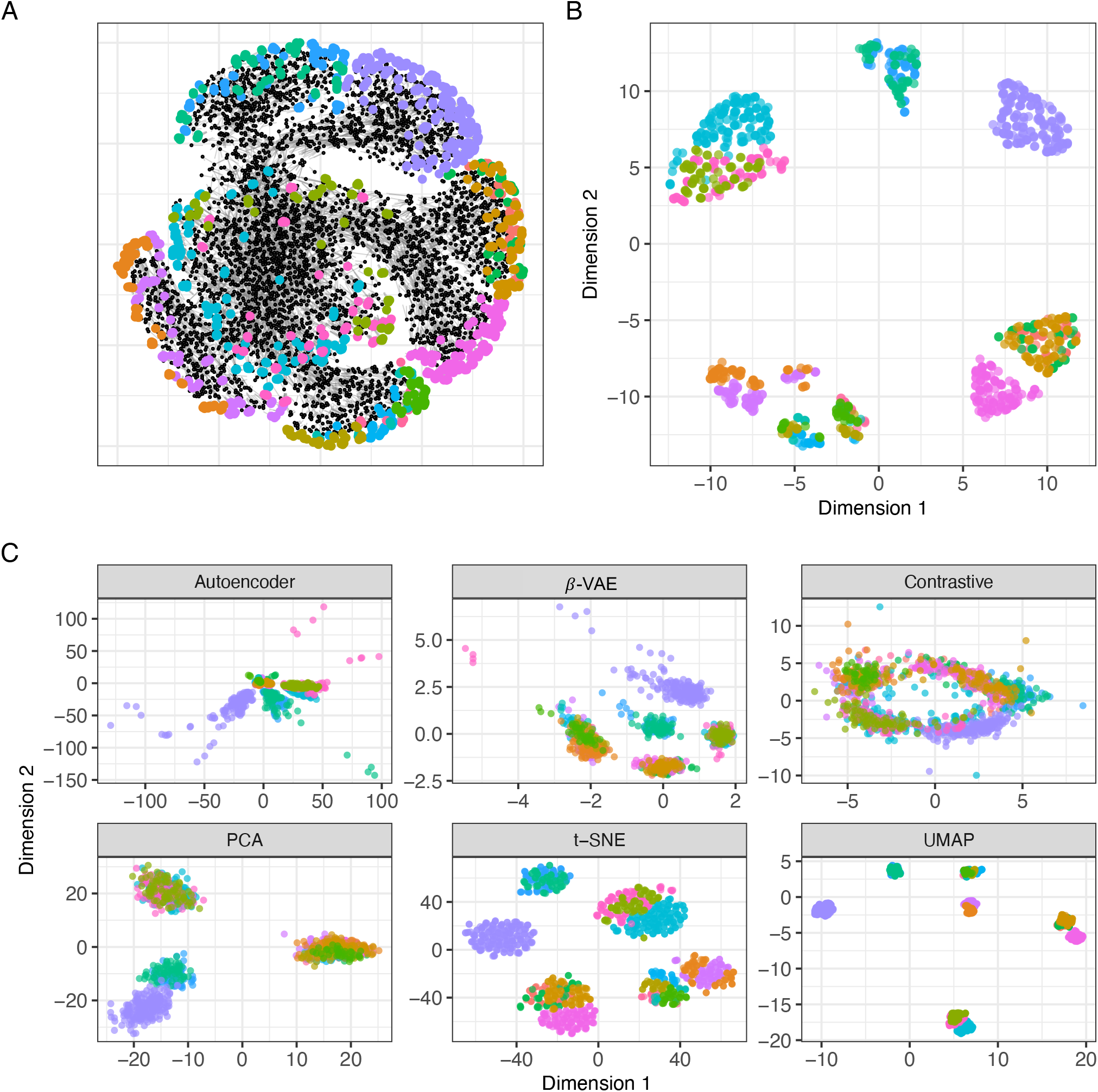
Using true genealogy (pedigree) as the truth to compare projection methods. In each plot, colors correspond to different sub-populations resulting from simulated migrations. (**A**) An example of a simulated population with migration and random mating showing ascendants (black nodes), genotyped descendants (colored nodes), and relationships (gray edges). (**B**) The “ground-truth” pairwise distances for the individuals of the last generation in all subpopulations, as calculated from the family tree, were used as the difference matrix in Multidimensional Scaling (MDS). This provides an example of what a visualization might look like if the distances between points represented differences in ancestry as accurately as possible. (**C**) Candidate visualizations of the simulated population.

### On the Success or Failure of Methods

It is important to note that the pedigree distance metric we develop here only measures how closely a visualization matches the viewer’s expectations of what it shows – that is, to what degree distances between points are indicative of differences in ancestry. The individual methods succeed or fail on this metric by incidence, not by design. For example, a method such as UMAP does not guarantee to preserve pairwise Euclidean distances between every pair of points, but rather, it seems to portray this incidentally in our experiments. Conversely, a method such as contrastive embedding learning is not a generally weak method for learning good representations of data – it only fails to portray what we expect an embedding method used in this domain to show.

### Limitations

Although we have attempted to quantify the “accuracy” of different visualizations with as much objectivity as possible, it remains true that visualization is inherently subjective. Even given these evaluations, practitioners may still prefer one visualization over another based on how it portrays certain known characteristics of the population. Therefore, our results only provide additional information for decision making based on a particular set of goals, not a blanket directive for which methods should be used.

All of the quantitative results reported here are associated with a particular set of assumptions. We find these assumptions to be reasonable – that distance between individuals should correlate with distance in ancestry – but not necessarily universal. For example, our evaluation metric assumes that this correlation should be linear (or linear following a log transform). Some researchers may prefer a method which exaggerates some types of distances, while minimizing others. This is not a problem, so long as the audience is aware of how the visualization should be interpreted. Other researchers may be either more or less tolerant to outliers than is reflected by our choice of Pearson’s *r* as the evaluation metric.

One weakness of the pedigree distance metric in natural datasets is that it does not measure the accuracy of distances between completely unrelated subpopulations, as there is no known pedigree relationship between these individuals. For example, a dataset containing two such subpopulations will attain the same performance on the pedigree distance metric whether the subpopulations are shown as completely separate or overlapping. This creates a possible bias in evaluation, as pedigree relationships are more likely to exist for closely related individuals – that is, the ground truth distances are more likely to exist between samples which reside in the same local neighbourhood. We have addressed this by using simulations, where ancestral relationships between every pair of individuals are known. When taking these distant relationships into account using the simulated data, we see that the same trends in performance hold for all of the methods, suggesting that this effect is likely not a significant drawback for our evaluations on the natural datasets.

We believe that our simulations (while not representative of every scenario) do capture general patterns of what we would expect in situations with selection and/or migration, as we show that the allele frequency spectrum shifts much more dramatically in the selected populations than in the randomly mated populations, regardless of migration (Figure 2). However, our simulations start out with uniform allele frequencies across sites, which may be simplistic but we feel captures the general trend of segregating sites in populations undergoing random and assortative mating.

**Figure 2.**
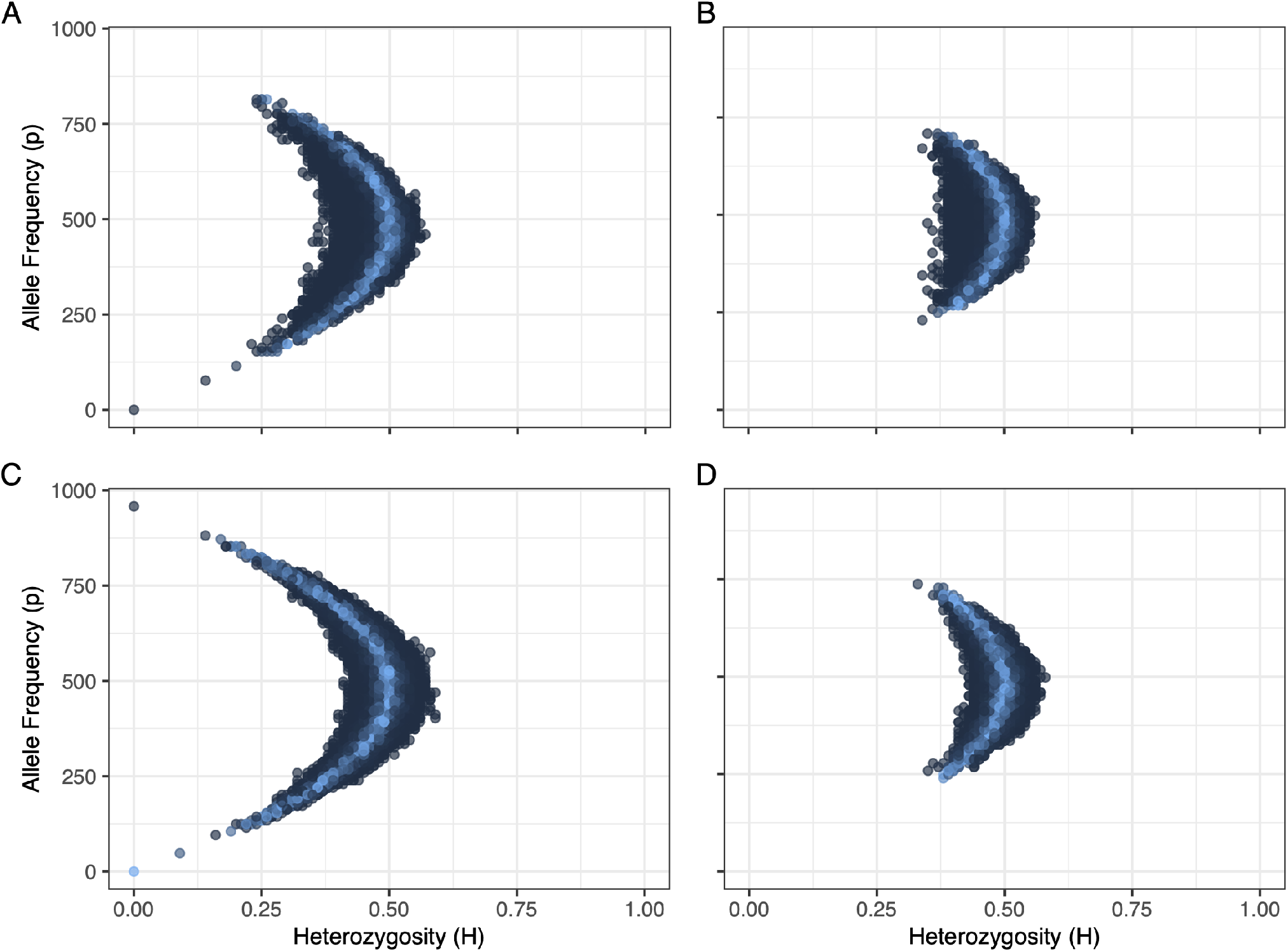
Allele frequency spectrum in a representative simulation with migration (**A, B**) and no migration (**C, D**) with assortative (**A, C**) and random (**B, D**) mating. Color represents the *χ*^2^ deviations from expected frequency with lighter colors representing larger p-values (less deviance).aim to reconstruct the graph in a lower-dimensional space, while maintaining the topological structure from the original high-dimensional space. This means that samples which are neighbours in the input space will tend to form neighbourhoods in the output space.

**Figure 3.**
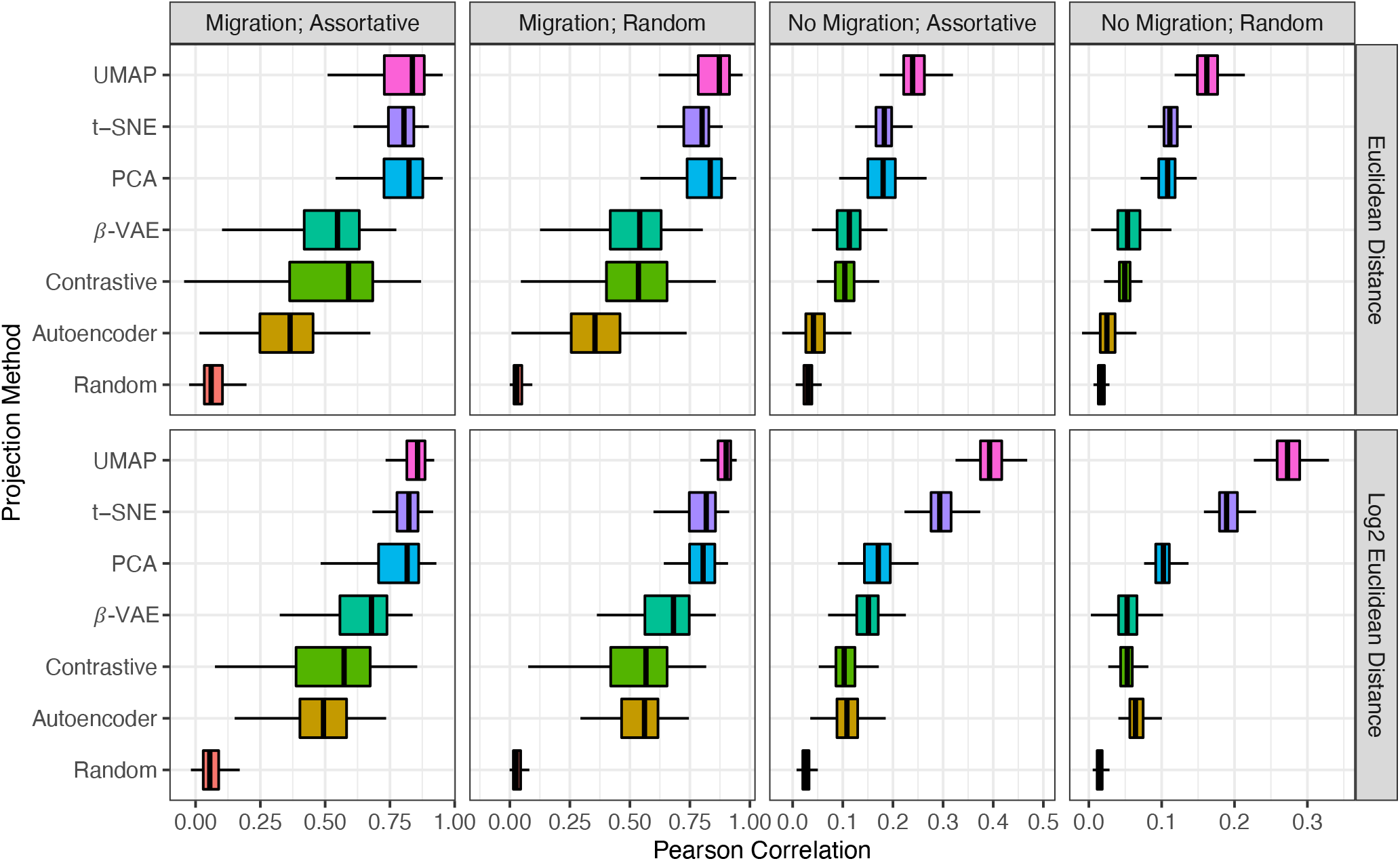
Correlation between the Euclidean distance (*D*; top row) and log_2_ of the Euclidean distance (log_2_(*D*); bottom row) in the visualization and the known pedigree distance in simulations with seven projection methods.

## Conclusions

To evaluate which embedding methods most accurately portray differences in ancestry, we have performed quantitative and qualitative analyses. Our analyses suggest that the best representation of genealogy, both globally and locally, are graph based algorithms such as t-SNE and UMAP. The results are consistent across real-world datasets as well as simulations, and for both random mating and assortative mating (breeding) populations with and without sub-population structure. Our results are also consistent whether evaluated on the basis of either linear or exponential distance with depth of ancestry. Our visualizations illustrate where nonlinear methods can improve on the embeddings provided by PCA, as well as how neural network based methods such as autoencoders and variational autoencoders fail in ways which are anticipated by theory.

## Materials & Methods

### Evaluation Using Pedigreed Populations

The objective evaluation of visualization methods is challenging. Given several options, practitioners may disagree on which visualization is subjectively the best for a given dataset.Battey et al. (2021) proposed to quantify the accuracy of a visualization by comparing the Euclidean distance between samples with their corresponding geographic distance. This is a good solution under the assumption that a correlation exists between geographic distance and distance in ancestry, which is generally true for any population where migration occurs. We extend this line of thinking by using pedigreed populations where ancestry can be calculated exactly, using the number of generations between ancestors and descendants. If ancestry is being represented well in the visualizations, then this distance should correlate with the Euclidean distance between the individuals in the embeddings. When the distance between ancestors and descendants is accurately modeled, the distance between, for example, siblings, is also enforced as they are anchored by their respective parents. In total, we use four publicly available datasets of genotyped individuals which include pedigree information.

#### Strawberry (Fragaria sp.)

We analyzed the global pedigree of wild and cultivated strawberry (*Fragaria sp.*) reported by Pincot et al. (2021). These pedigree records were assembled for 8,851 individuals, including 2,656 cultivars developed since 1775. SNP marker genotypes for 1,495 individuals were available, including 1,235 UCD and 260 USDA accessions (asexually propagated individuals) previously geno-typed byHardigan et al. (2018) with the iStraw35 SNP array (Bassil et al., 2015; Verma et al., 2016). This pedigree extends 23 generations for 2 ascendant-descendent pairs and has a median depth of 7 generations. The strawberry dataset included both natural accessions and elite Californian breeding material; the majority of samples are from the latter and the former includes diverse samples from Asia, Europe, North America, and South America. Despite this,Pincot et al. (2021) suggests a high level of admixture and allele sharing among the individuals in the metropolitan population.

#### Pig (Sus scrofa)

We analyzed a pig dataset that PIC (a Genus company) made available for comparing genomic prediction methods by Cleveland et al. (2012). The dataset contains 3,534 individuals with 52,842 SNP genotypes from the Illumina PorcineSNP60 chip (Ramos et al., 2009) and a pedigree including parents and grandparents of the genotyped animals (Cleveland et al., 2012). This pedigree extends 14 generations for 417 ascendant-descendant pairs and has a median depth of 7 generations. The PIC Pig population is a strictly a breeding population and we expect it to be without substantial population structure or well defined sub-populations.

#### Soay Sheep (Ovis aries)

We analyzed the wild Soay sheep population reported by Stoffel et al. (2021). This dataset includes genetypes from the Illumina Ovine SNP50 BeadChip containing 51,135 SNP markers for 5,952 wild Soay sheep with both genotypes and pedigree. There are 8,172 entries in the pedigree records. This pedigree extends 13 generations for 1 ascendant-descendant pair and has a median depth of 5 generations. The Soay Sheep is a natural population in which ascendants and descendants are genotypes and pedigreed and we expect there to be limited population structure or well defined sub-populations.

#### Florida Scrub-Jay (Aphelocoma coerulescens)

We analyzed a natural Florida Scrub-Jay population from Chen et al. (2019). Our final pedigree consists of 6,936 individuals with truncated birth years post 1990 with SNP genotypes of 10,731 autosomal SNPs from 3,404 individuals (Chen et al., 2019). This pedigree extends 8 generations for 54 ascendant-descendent pairs and has a median depth of 2 generations. The Florida Scrub-Jay population is a wild population including genotypes for ascendants and descendants and we expect there to be limited population structure or well defined subpopulations.Chen et al. (2019) notes that the genetic contribution of immigrants to be 75% in the final population, which suggests a high degree of admixture.

### Evaluation Using Simulation

In order to provide a complete ground truth for evaluation, which includes known pairwise ancestral distances between both distantly and immediately related individuals, we use a variety of population-level simulations. These simulations are based on the SeqBreed software package (Pérez-Enciso et al., 2020), which was modified to begin from a hypothetical founder population with a uniformly random distribution of alleles. This ensures that all of the population structure present in the data is from the simulation. An example of one such simulated population is shown in Figure 1.

Each simulation started from a diploid founding population of one hundred individuals, and proceeded for ten generations. A constant recombination rate of 1cM/Mbp is assumed, and no sex chromosomes were simulated. In total, 24,000 biallelic SNPs were simulated in each population and were used to genotype the terminal nodes (extant individuals).

We performed simulations with migration, and without migration. For the simulations with migration, half of the current generation emigrates to found a separate subpopulation with some probability (*p* = 0.3). For each of the simulations with and without migration, we performed versions with random and assortative mating.

For the simulations with random mating, every individual within the current generation pairs with a random other individual. A random number of offspring, between one and four, are produced. This resulted in an average *F_ST_* = 0.083, 0.24 ≤ *MAF* ≤ 0.5 which is concordant with low genetic differentiation between the known subpopulations, expected in a stable, random mating population.

For simulations with assortative mating, we defined a hypothetical quantitative trait with 10 randomly selected QTN (*h*^2^ = 0.5). For each generation, the top 50% of the individuals are selected based on their phenotype, and each pairing produces between two and eight offspring. This resulted in an average *F_ST_* = 0.155, 0 ≤ *MAF* ≤ 0.5 which is concordant with moderate genetic differentiation between the known subpopulations. The populations are twice as differentiated. Simulations were run 100 times each to obtain error estimates. Population genetic parameters are estimated using *popgen(*) from snpReady v0.9.6 (Granato and Fritsche-Neto, 2018) in R v4.1.0 (R Core Team, 2021). The allele frequency spectrum as a function of heterozysosity for each site is shown in Figure 2 (Ferretti et al., 2018).

### Models Evaluated

For all evaluations, we compare PCA, t-SNE, UMAP, an autoencoder, and a *β*-VAE (Higgins et al., 2017). In addition, we use a learned embedding obtained via an unsupervised contrastive technique (Ye et al., 2019). We also use a random nonlinear projection from a randomly initialized two-layer neural network as a comparison baseline. Since, in practice, the ancestral relationships are unknown, it is not realistic to tune each method specifically to maximize performance on the pedigree distance task. For the neural network based methods, we choose hyperparameters which allow the training loss to decrease to a plateau. For the Autoencoder, *β*-VAE, and contrastive methods, we choose a standard encoder and decoder architecture.

When using techniques such as t-SNE and UMAP, it is common to preprocess the data using PCA and use the first *n* principal components as the input. However, we found that for the pedigree distance metric, the raw marker data provided better performance. Both methods are equipped with hyperparameters which can be adjusted to change the characteristics of the resulting visualization. In the interest of fairness, we use only the defaults for each. We use the umap-learn implementation for UMAP (McInnes et al., 2018) and the scikit-learn implementation for t-SNE (Pedregosa et al., 2011).

When testing the VAE with the configuration proposed in Battey et al. (2021) (which we will refer to as *popvae*), we observed a strong propensity for posterior collapse. This is a common failure mode for VAEs, especially those with powerful decoders as in popvae. Posterior collapse is signalled by the posterior of the model collapsing to the prior, such that **z** contains no information about **x**. Many authors have explored the problem of posterior collapse, and have suggested solutions such as an annealing schedule for the KL term, bounds on log-likelihood, or hierarchical latent dependencies (Dai et al., 2020). One of the simplest solutions for mitigating posterior collapse is a gradual warm-up where a weight on the KL term is initialized at zero and increased according to an annealing schedule until it reaches one (Bowman et al., 2015). We found that training was unstable with either monotonic or cyclical annealing schedules, with the KL loss term often exploding to an unrecoverable state. However, adding a constant coefficient of *β* = 0.001 to the KL term was effective in preventing posterior collapse and did not require hand-tuning on a per-dataset basis. This simple modification is known as the *β*-VAE and was originally proposed to encourage disentanglement between latent factors in VAEs (Higgins et al., 2017). Setting *β* = 1 recovers the vanilla VAE, while *β* > 1 emphasizes disentanglement of the factors of variation by increasing the variational bottleneck. However, setting *β* < 1 as we have done here is protective against posterior collapse by simply relaxing the regularization imposed by the KL term. This significantly damages the model’s ability to generate unconditional samples (Hoffman et al., 2017) – however, here we are only interested in the inference model.

In general, we found the behavior of the VAE to be very different in each dataset. Relaxing the KL term to prevent collapse was not necessary for every dataset, and can in fact have a small negative effect on some datasets – however, in the interest of presenting methods which are not hand-tuned for each individual dataset, we use it across datasets regardless. In Battey et al. (2021), the authors describe an issue which they refer to as overfitting. The authors propose to remedy this problem by using early stopping based on held-out samples. We found this to be the case for some datasets when using the *β*-VAE, because the relaxation of the KL term means that outliers can move away from the clusters into areas of extremely low density under the prior. We found early stopping to be less important for the vanilla VAE, however, with some datasets the performance did show small fluctuations over the number of training epochs. We decided to not include early stopping in our experiments, as holding out data for early stopping also means that it is impossible to compare the method to others as it does not take into account all of the data.

Unsupervised embedding learning is a commonly used technique for performing tasks such as information retrieval in unlabeled datasets. These techniques attempt to learn an embedding space where similar samples lie close together in the space and dissimilar samples are farther apart. For our purposes, we chose to use a contrastive embedding learning technique with instance-wise supervision (Ye et al., 2019). Training samples **x** are augmented to create positive samples 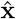. This is done by randomly changing homozygous sites to heterozygous with some probability (*p* = 0.1). We then minimize the loss function

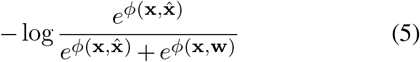

where *ϕ* is a similarity metric, here we use the cosine similarity, and **w** is a different sample than **x**. We found that cosine similarity performs better on the pedigree distance metric than other similarity metrics, such as inverse Euclidean distance.

## References

Arvanitidis, G., Hansen, L. K., and Hauberg, S. (2017). Latent space oddity: on the curvature of deep generative models. arXiv preprint arXiv:1710.11379.

Ausmees, K. and Nettelblad, C. (2020). A deep learning framework for characterization of genotype data. bioRxiv.

Bassil, N. V., Davis, T. M., Zhang, H., Ficklin, S., Mittmann, M., Webster, T., Mahoney, L., Wood, D., Alperin, E. S., Rosyara, U. R., et al. (2015). Development and preliminary evaluation of a 90 k axiom® snp array for the allooctoploid cultivated strawberry fragaria×ananassa. BMC genomics, 16(1):1–30.

Battey, C. J., Coffing, G. C., and Kern, A. D. (2021). Visualizing population structure with variational autoencoders. G3 Genes|Genomes|Genetics, 11(1).

Bowman, S. R., Vilnis, L., Vinyals, O., Dai, A. M., Jozefowicz, R., and Bengio, S. (2015). Generating sentences from a continuous space. arXiv preprint arXiv:1511.06349.

Chen, N., Juric, I., Cosgrove, E. J., Bowman, R., Fitzpatrick, J. W., Schoech, S. J., Clark, A. G., and Coop, G. (2019). Allele frequency dynamics in a pedigreed natural population. Proceedings of the National Academy of Sciences, 116(6):2158–2164.

Cleveland, M. A., Hickey, J. M., and Forni, S. (2012). A common dataset for genomic analysis of livestock populations. G3: Genes| Genomes| Genetics, 2(4):429–435.

Dai, B., Wang, Z., and Wipf, D. (2020). The usual suspects? reassessing blame for vae posterior collapse. In International Conference on Machine Learning, pages 2313–2322. PMLR.

Diaz-Papkovich, A., Anderson-Trocmé, L., Ben-Eghan, C., and Gravel, S. (2019). UMAP reveals cryptic population structure and phenotype heterogeneity in large genomic cohorts. PLoS Genetics, 15(11):1–24.

Ferretti, L., Ribeca, P., and Ramos-Onsins, S. E. (2018). The site frequency/dosage spectrum of autopolyploid populations. Frontiers in genetics, 9:480.

Francis, R. M. (2017). pophelper: an r package and web app to analyse and visualize population structure. Molecular ecology resources, 17(1):27–32.

Granato, I. and Fritsche-Neto, R. (2018). snpReady: Preparing Genotypic Datasets in Order to Run Genomic Analysis. R package version 0.9.6.

Hardigan, M. A., Poorten, T. J., Acharya, C. B., Cole, G. S., Hummer, K. E., Bassil, N., Edger, P. P., and Knapp, S. J. (2018). Domestication of temperate and coastal hybrids with distinct ancestral gene selection in octoploid strawberry. The plant genome, 11(3):180049.

Higgins, I., Matthey, L., Pal, A., Burgess, C., Glorot, X., Botvinick, M., Mohamed, S., and Lerchner, A. (2017). beta-vae: Learning basic visual concepts with a constrained variational framework. In Proc. ICLR.

Hoffman, M. D., Riquelme, C., and Johnson, M. J. (2017). The *β*-vae’s implicit prior. In NIPS Workshop on Bayesian Deep Learning.

Holsinger, K. E. and Weir, B. S. (2009). Genetics in geographically structured populations: defining, estimating and interpreting f st. Nature Reviews Genetics, 10(9):639–650.

Huszár, F. (2017). Variational inference using implicit distributions. arXiv preprint arXiv:1702.08235.

Kingma, D. P. and Welling, M. (2013). Auto-encoding variational bayes. arXiv preprint arXiv:1312.6114.

Li, W., Cerise, J. E., Yang, Y., and Han, H. (2017). Application of t-sne to human genetic data. Journal of Bioinformatics and Computational Biology, 15(04):1750017.

López-Cortés, X. A., Matamala, F., Maldonado, C., Mora-Poblete, F., and Scapim, C. A. (2020). A Deep Learning Approach to Population Structure Inference in Inbred Lines of Maize. Frontiers in Genetics, 11(November):1–8.

Marnetto, D. and Huerta-Sánchez, E. (2017). Haplostrips: revealing population structure through haplotype visualization. Methods in Ecology and Evolution, 8(10):1389–1392.

McInnes, L., Healy, J., Saul, N., and Grossberger, L. (2018). Umap: Uniform manifold approximation and projection. The Journal of Open Source Software, 3(29):861.

Meisner, J. and Albrechtsen, A. (2020). Haplotype and population structure inference using neural networks in wholegenome sequencing data. bioRxiv.

Patterson, N., Price, A. L., and Reich, D. (2006). Population structure and eigenanalysis. PLoS genetics, 2(12):e190.

Pedregosa, F., Varoquaux, G., Gramfort, A., Michel, V., Thirion, B., Grisel, O., Blondel, M., Prettenhofer, P., Weiss, R., Dubourg, V., Vanderplas, J., Passos, A., Cournapeau, D., Brucher, M., Perrot, M., and Duchesnay, E. (2011). Scikit-learn: Machine learning in Python. Journal of Machine Learning Research, 12:2825–2830.

Pérez-Enciso, M., Ramírez-Ayala, L. C., and Zingaretti, L. M. (2020). SeqBreed: A python tool to evaluate genomic prediction in complex scenarios. Genetics Selection Evolution, 52(1):1–9.

Petkova, D., Novembre, J., and Stephens, M. (2016). Visualizing spatial population structure with estimated effective migration surfaces. Nature genetics, 48(1):94–100.

Pincot, D. D., Ledda, M., Feldmann, M. J., Hardigan, M. A., Poorten, T. J., Runcie, D. E., Heffelfinger, C., Dellaporta, S. L., Cole, G. S., and Knapp, S. J. (2021). Social network analysis of the genealogy of strawberry: retracing the wild roots of heirloom and modern cultivars. G3, 11(3):jkab015.

Platzer, A. (2013). Visualization of SNPs with t-SNE. PLoS ONE, 8(2).

Price, A. L., Patterson, N. J., Plenge, R. M., Weinblatt, M. E., Shadick, N. A., and Reich, D. (2006). Principal components analysis corrects for stratification in genome-wide association studies. Nature genetics, 38(8):904–909.

R Core Team (2021). R: A Language and Environment for Statistical Computing. R Foundation for Statistical Computing, Vienna, Austria.

Ramos, A. M., Crooijmans, R. P., Affara, N. A., Amaral, A. J., Archibald, A. L., Beever, J. E., Bendixen, C., Churcher, C., Clark, R., Dehais, P., et al. (2009). Design of a high density snp genotyping assay in the pig using snps identified and characterized by next generation sequencing technology. PloS one, 4(8):e6524.

Steinig, E. J., Neuditschko, M., Khatkar, M. S., Raadsma, H. W., and Zenger, K. R. (2016). netview p: a network visualization tool to unravel complex population structure using genome-wide snp s. Molecular Ecology Resources, 16(1):216–227.

Stoffel, M. A., Johnston, S. E., Pilkington, J. G., and Pemberton, J. M. (2021). Genetic architecture and lifetime dynamics of inbreeding depression in a wild mammal. Nature Communications, 12(1):1–10.

Van Heerwaarden, J., Doebley, J., Briggs, W. H., Glaubitz, J. C., Goodman, M. M., Gonzalez, J. d. J. S., and Ross-Ibarra, J. (2011). Genetic signals of origin, spread, and introgression in a large sample of maize landraces. Proceedings of the National Academy of Sciences, 108(3):1088–1092.

Verma, S., Bassil, N., Van De Weg, E., Harrison, R., Monfort, A., Hidalgo, J., Amaya, I., Denoyes, B., Mahoney, L., Davis, T., et al. (2016). Development and evaluation of the axiom® istraw35 384ht array for the allo-octoploid cultivated strawberry fragaria×ananassa. In VIII International Strawberry Symposium 1156, pages 75–82.

Vincent, P., Larochelle, H., Bengio, Y., and Manzagol, P.-A. (2008). Extracting and composing robust features with denoising autoencoders. In Proceedings of the 25th international conference on Machine learning, pages 1096–1103.

Ye, M., Zhang, X., Yuen, P. C., and Chang, S.-F. (2019). Unsupervised embedding learning via invariant and spreading instance feature. In Proceedings of the IEEE/CVF Conference on Computer Vision and Pattern Recognition, pages 6210–6219.

Yu, J., Pressoir, G., Briggs, W. H., Bi, I. V., Yamasaki, M., Doebley, J. F., McMullen, M. D., Gaut, B. S., Nielsen, D. M., Holland, J. B., et al. (2006). A unified mixed-model method for association mapping that accounts for multiple levels of relatedness. Nature genetics, 38(2):203–208.

